# Does body position before and during blood sampling influence the athlete biological passport variables?

**DOI:** 10.1101/759563

**Authors:** Astolfi Tiffany, Schumacher Yorck Olaf, Crettaz von Roten Fabienne, Saugy Martial, Faiss Raphael

**Author notes:** Corresponding author: Raphael Faiss, Center of Research and Expertise in anti-Doping sciences - REDs, University of Lausanne, Switzerland.

## Abstract

The Athlete’s Biological Passport (ABP) is a tool for the indirect detection of blood doping. Current guidelines from the World Anti-Doping Agency (WADA) require a delay of 2 hours after any physical exercise and to be seated for 10 minutes prior to any blood sampling to obtain a valid measurement. Since body position prior to and during phlebotomy may influence the outcome, this study compared blood biomarker variations with changes in body position during blood sample collection. Ten successive venous blood samples from 38 subjects of 3 groups (elite cyclists, apnea divers and controls) in three situations (seated, after a 50 m walk, and supine) were collected and analyzed via flow cytometry. While reticulocytes percentage was unchanged in all conditions, haemoglobin concentration and hematocrit were stable after at least 10 min in a seated position. Due to shifts in plasma volume, the measures were significantly higher after changing posture for a short walk, but readjusted to previous levels after only 5 min. Supine position caused generally lower values after 10-30 min. The results support the current guidelines and additionally provide evidence to adjust the waiting time for blood sampling after short changes in posture.

## Introduction

Nearly 30000 blood samples are collected yearly for the athlete biological passport (ABP). The ABP tracks blood markers longitudinally and based on changes in these variables, helps identifying patterns of doping (Sottas, Robinson et al. 2011, Vernec 2014). However, despite remarkable results since its implementation, athletes are getting used to fine-tuning doping methods to circumvent the ABP testing strategies. Actually, it is now certain that athletes adapted their protocols using rather first generation of rhEPO and frequent micro-dosing in the evening to maintain a supraphysiological level of haemoglobin while reducing the chances of being tested positive (Hamilton and Coyle 2012). Therefore, accurate and precise measurement of blood variables with low bias are paramount to ensure the indirect detection and targeting potential of the ABP. The blood matrix as a suspension of living cells in plasma transports oxygen to the working muscles. Variations in the fluid balance and thus plasma volume may inevitably alter variables for which concentration values are reported (e.g., haemoglobin concentration ([Hb]) or haematocrit (Hct)) while absolute measures (e.g., reticulocytes percentage (Ret%)) remain stable (Fahraeus 1929, Ahlgrim, Pottgiesser et al. 2010). Significant shifts in plasma volume were hence reported in connection with many situations of an athletes’ daily life such as acute physical exercise, heat exposure, psychological stress and/or postural changes (Harrison 1975, Harrison and Edwards 1976, Collins, Hill et al. 1986, Imelik and Mustimets 1992, Lundvall and Bjerkhoel 1995, Bloomer and Farney 2013). Acute or chronic plasmatic volume variations may also occur with endurance training periods (Sawka, Convertino et al. 2000), repeated sauna bathing (Stanley, Halliday et al. 2015), during altitude acclimatization (Sawka, Young et al. 1996) or hypoxic exposure with a high individual variability (Young, Karl et al. 2019). Overall, this underlines the numerous confounding factors affecting concentration based blood markers and the need for robust procedures to limit pre-analytical variations when analysed for the ABP. For instance, the World Anti-Doping Agency (WADA) enacted specific blood collection guidelines (WADA 2016) in addition to precise Blood Analytical Requirements for the Athlete Biological Passport (WADA 2009). Currently, the guidelines specify that 2 h waiting is necessary after any physical exercise and require the athletes to be seated for 10 min before sample collection to allow the vascular volumes to equilibrate. Effects of posture upon blood volume and composition have been extensively investigated (Thompson, Thompson et al. 1928). Recent updates highlight the need for standardization because position changes may rapidly alter plasma volume (Lippi, Salvagno et al. 2015, Lima-Oliveira, Guidi et al. 2017). In an antidoping context, a prior study investigated if 10 min of seating was enough to guarantee stable [Hb] and Hct readings for the ABP (Ahlgrim, Pottgiesser et al. 2010).

In a more general context, the world health organizations’ current best practice for blood drawing recommends to “make the patient comfortable in a supine position (if possible)” (World Health Organisation 2010). The latter is however, not the standard for blood collection in an antidoping context but might be more comfortable for certain athletes.

The aim of this study was thus to investigate the influence of body position prior to and during phlebotomy in this context (i.e. seated vs. supine). To improve practical application, this study assessed if a short position change (e.g., walking a short distance from a waiting room to the sample collection site) influences the readings and may thus be acceptable in the context of the normal antidoping blood sample collection sequence.

## Material and methods

### Study subjects

Thirty-eight non-smoking healthy Caucasian subjects were included in this study in three groups: 10 elite cyclists (Cyc, 10 males), 12 trained apnea divers (Apn, 7 males, 5 females) and 16 moderately trained control subjects (Con, sport sciences students, 9 males, 7 females). The Cyc group included International Elite licensed cyclists successful at an international level (e.g., UCI World Tour races or UCI World Cup participations) in road, track, and mountain-bike events. The Apn group included apnea divers with a regular practice of apnea training (i.e. at least two weekly training sessions with or without immersion), competing at a national or international level (breath holding experience 8 ± 2.3 yrs). Inclusion criteria for the control group was a total weekly volume of aerobic sport activities (e.g., running, triathlon, ski-cross country) not exceeding 4 hours. Participants living permanently at an altitude above 800 m were also excluded. The procedure and risks were fully explained to the subjects, and all of them gave their written consents to participate in this study. This study was approved by the local ethics committee (CCER-VD, Lausanne, Switzerland, Agreement 2018-01019) and conducted in respect of the Declaration of Helsinki.

### Study design, pre-analytical conditions and hematological analyses

To mimic as closely as possible an ABP blood sample collection, the current WADA guidelines on analytical procedures were strictly followed (WADA 2016). A total of 10 successive blood samples were collected from each subject over a single visit of 70 minutes in the same sequence. The study design is illustrated in Figure 1. At least 2 hours after any physical activity or exercise training, subjects reported to the laboratory as part of their daily activity. Room temperature was kept constant in the laboratory at approx. 21° C.

**Figure 1.**
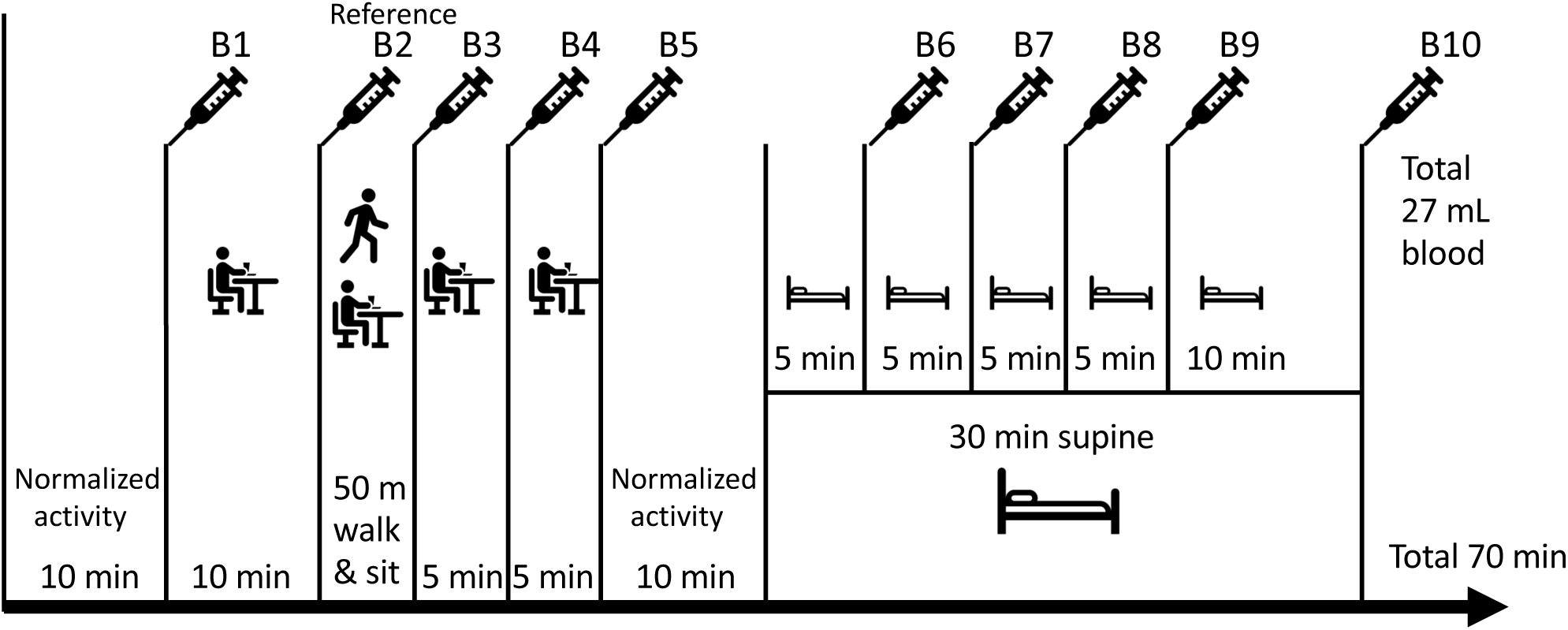
Study design and blood collection times seated or supine

A 10-min period of normalized activity was imposed upon arrival to the laboratory: walking 1 min, sitting down to read a newspaper (4 min), walking down the stairs one floor and up again (1 min), waiting in a standing position (2 min) before walking to the phlebotomy lab (2 min). Subjects were then asked to sit down and a first blood sample was taken within 1 min (B1). Subsequently, subjects remained seated for 10 min and were requested to fill in a food and exercise training diary for the last 24 hours prior to the laboratory visit to control for their activity and hydration status. A second blood sample was taken after 10 min seated (B2). In an antidoping context, this sample typically corresponds to a reference sample that could have been collected for an ABP analysis. Subjects were then asked to stand up and walk exactly 50 m before a third sample was drawn in order to mimic a situation where an athlete in a waiting room would have to change seat for the blood collection (B3). After 5 and 10 minutes in the seated position, two other samples were drawn (B4 & B5). Immediately after, subjects were requested to replicate the initial 10-min period of standardized activity before lying down on an examination table and remaining in a supine position with the head slightly above the rest of the body (back of the examination table angle of 30°). Blood samples were then taken exactly after 5, 10, 15, 20 and 30 minutes in the supine position (B6-B10).

“Butterfly” 21G needles with a short manifold were inserted in one of the antecubital veins (Sarstedt Safety-Multifly^®^, Sarstedt AG, Nuembrecht, Germany) after proper disinfection of the site. Intravenous access using a butterfly was preferred to reduce hemolysis (Barnaby, Wollowitz et al. 2016). A tourniquet was used as standard to facilitate puncture and removed once the butterfly was inserted in the antecubital vein and fixed with medical tape. Tourniquet time was approx. 25 s and never exceeded 60 s as required by the WADA guidelines (WADA 2016). Blood was collected in two EDTA-coated tubes (Sarstedt S-Monovette 2.7 ml, Sarstedt AG, Nuernbrecht, Germany). The first tube served to purge the butterfly manifold and was thrown. Tubes were inverted 10 times directly after blood collection in order to ensure proper mixing with the anti-coagulant. All blood samples were collected by the same experienced phlebotomist throughout the study. In the case of a blood clot in the bsutterfly manifold, another puncture on another site was made. Four samples were collected in the seated position from one arm and the subsequent 6 blood samples in the supine position from the other arm. The first arm was chosen randomly.

Blood variables were determined via flow cytometry using a fully automated cell counter (Sysmex XN1000, Sysmex Europe GmbH, Norderstedt, Germany). Internal quality controls provided by the manufacturer) were run before each analytical batch as described thoroughly elsewhere (Robinson, Saugy et al. 2018). Once collected, blood samples were stored in a fridge at 4° C for a time lasting between 30 minutes and 12 hours. All samples from one single subject were subsequently analyzed at the same time as part of the same batch after being rolled during 15 minutes at room temperature for homogenization and temperature stabilization purposes. All samples were analyzed at least twice in order to ensure valid recording of [Hb] and Ret% according to the WADA guideline in force (WADA 2019).

At the end of the blood sampling procedure, circulating volumes (e.g. blood volume, plasma volume, and total hemoglobin mass (Hbmass)) was measured with a carbon monoxide (CO) rebreathing technique with a fully automated system (OpCo: Detalo Instruments, Birkerod, Denmark) as described elsewhere (Siebenmann, Keiser et al. 2017)

Plasma volume changes (ΔPV) were calculated using the formula introduced by Dill and Costill (Costill and Fink 1974): 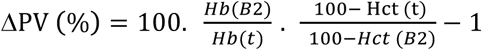

[Hb] and Hct measured at B2 (i.e. after 10 min seated) were used as reference baseline values; t indicates the different collection timepoints (from B1 to B10).

Blood pressure was additionally monitored with an automated wrist manometer (Beurer BC85, Beurer Gmbh, Ulm, Germany) at B1 and B6 to verify that the normalized activity imposed for the protocol only produced a low sympathetic stimulus.

### Statistical analyses

Descriptive values are reported as means ± SD. Variability of [Hb] and Hct over time was calculated as a coefficient of variation (CV, %) from the mean of each individuals’ CV over the 10 measurements. Normality of the distributions was successfully tested with the Shapiro-Wilk test. Sphericity was not assumed and the Geisser-Greenhouse correction was used. Baseline variables were compared between groups with Students’ t test. Differences in [Hb], Hct and Ret% at the different time points were thus assessed with a linear mixed-model procedure with fixed and random effects to explain target variables where subjects represented random effects; time, sex and group were the fixed effects. The F statistic was used to test for significant fixed effects. The repeated measures were analyzed by comparing each time point with the reference value (at B2) with correction for multiple comparisons using statistical hypothesis testing (Dunnett’s test). The null hypothesis was rejected for *P*<0.05. All statistical analyses were performed using the Jamovi open-source dedicated statistical software (Jamovi 2019).

## Results

Anthropometrical data, variables measured at baseline (or after all blood samples for total haemoglobin mass and circulating volumes), for the study subjects are reported in Table 1. Hbmass (absolute and relative to body mass) and circulating volumes were significantly higher in the Cyc group when compared to Apn and Con (Table 1). In females, Hbmass (636 ± 84 g) and blood volume (4864 ± 646 ml) were significantly lower than in males (957 ± 121 g and 6748 ± 1054 ml, respectively, *P*<0.001). Similarly, in females, [Hb] (12.7 ± 0.7 g/dL) and Hct (37 ± 2.1%) were significantly lower (*P*<0.01) than in males (14.3 ± 0.9 g/dL) for [Hb], and (40.8 ± 2.2%) for Hct.

**Table 1.** Anthropometrical data and baseline variables (at B2), otherwise specified. Mean values ± SD.

|  | Cyclists<br>(n=10) | Apnea divers<br>(n=12) | Controls<br>(n=16) |
| --- | --- | --- | --- |
| Males/Females | 10/0 | 7/5 | 9/7 |
| Age (years) | 28.4 ± 6 | 37.0 ± 9*<sup>##</sup> | 24.8 ± 3 |
| Height (cm) | 180 ± 6 | 176 ± 9 | 173 ± 7** |
| Body mass (kg) | 72.2 ± 7.0 | 71.9 ± 11.2 | 68.3 ± 10.0 |
| Total haemoglobin mass (g) | 988 ± 94 | 863 ± 189* | 820 ± 208* |
| Total haemoglobin mass (g/kg) | 14.0 ± 0.4 | 11.9 ± 1.9* | 11.5 ± 1.8** |
| Plasma volume (ml) | 4338 ± 598 | 3638 ± 727* | 3888 ± 598* |
| Blood volume (ml) | 7201 ± 789 | 6093 ± 1197* | 5764 ± 1348* |
| Haemoglobin concentration (g/dL) | 13.8 ± 0.6 | 13.9 ± 1 | 13.7 ± 1.3 |
| Hematocrit (%) | 39.9 ± 1.9 | 39.8 ± 2.1 | 39.5 ± 3.0 |
| Systolic Blood pressure at B1 (mmHg) | 124 ± 10 | 118 ± 9 | 118 ± 8 |
| Diastolic Blood pressure at B1 (mmHg) | 70 ± 8 | 79.3 ± 6* | 74.5 ± 10 |
| Systolic Blood pressure at B6 (mmHg) | 128 ± 13 | 127 ± 4 | 126 ± 16 |
| Diastolic Blood pressure at B6 (mmHg) | 76 ± 7 | 82.8 ± 8 | 75 ± 7 |

\*\* $P < 0.01$ , \* $P < 0.05$ for difference with Cyclists
<sup>##</sup> $P < 0.01$ for difference with Controls.

Box-and-whisker plots for [Hb], Hct, and Ret% over the 10 timepoints are presented in Figure 2. Average coefficients of variation for all subjects over all time points for [Hb] and Hct were of 1.4 ± 0.2% for [Hb] and 1.6 ± 0.4% for Hct.

**Figure 2.**
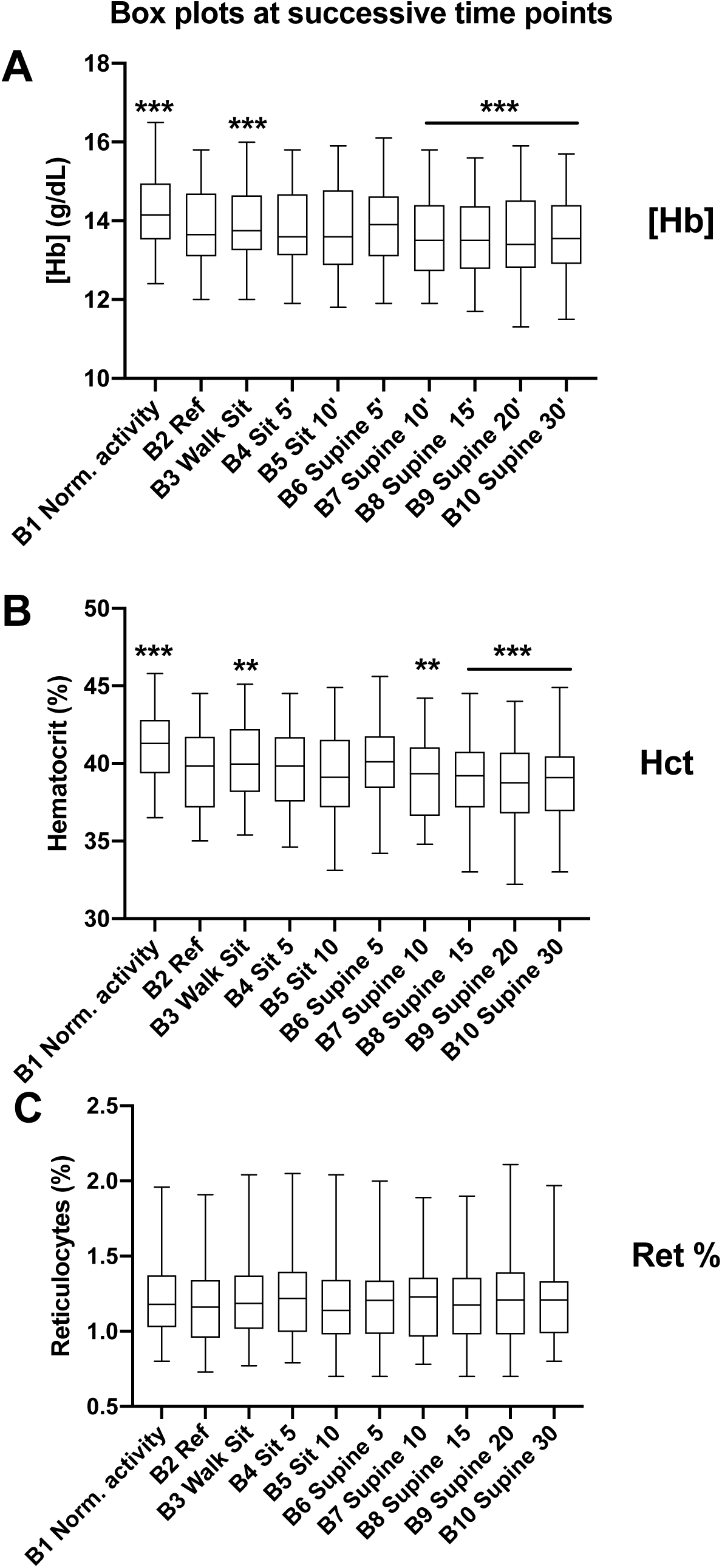
Box plots at successive timepoints (B1 to B10) for hemoglobin concentration (A), hematocrit (B), and reticulocytes % (C) for all subjects (n=38). *** *P*<0.001, ** *P*<0.01 for difference with baseline (B2 Ref)

When compared to the B2 reference value, significant differences were observed after the initial normalized activity (B1), after 50 m of walking (B3) and in the supine position (B7-B10) for [Hb] and Hct but not for Ret% (Figure 3). No significant difference was observed in the reticulocytes percentage (Ret %) over the 10 timepoints (F (9, 324) =1.13, P = 0.339).

**Figure 3.**
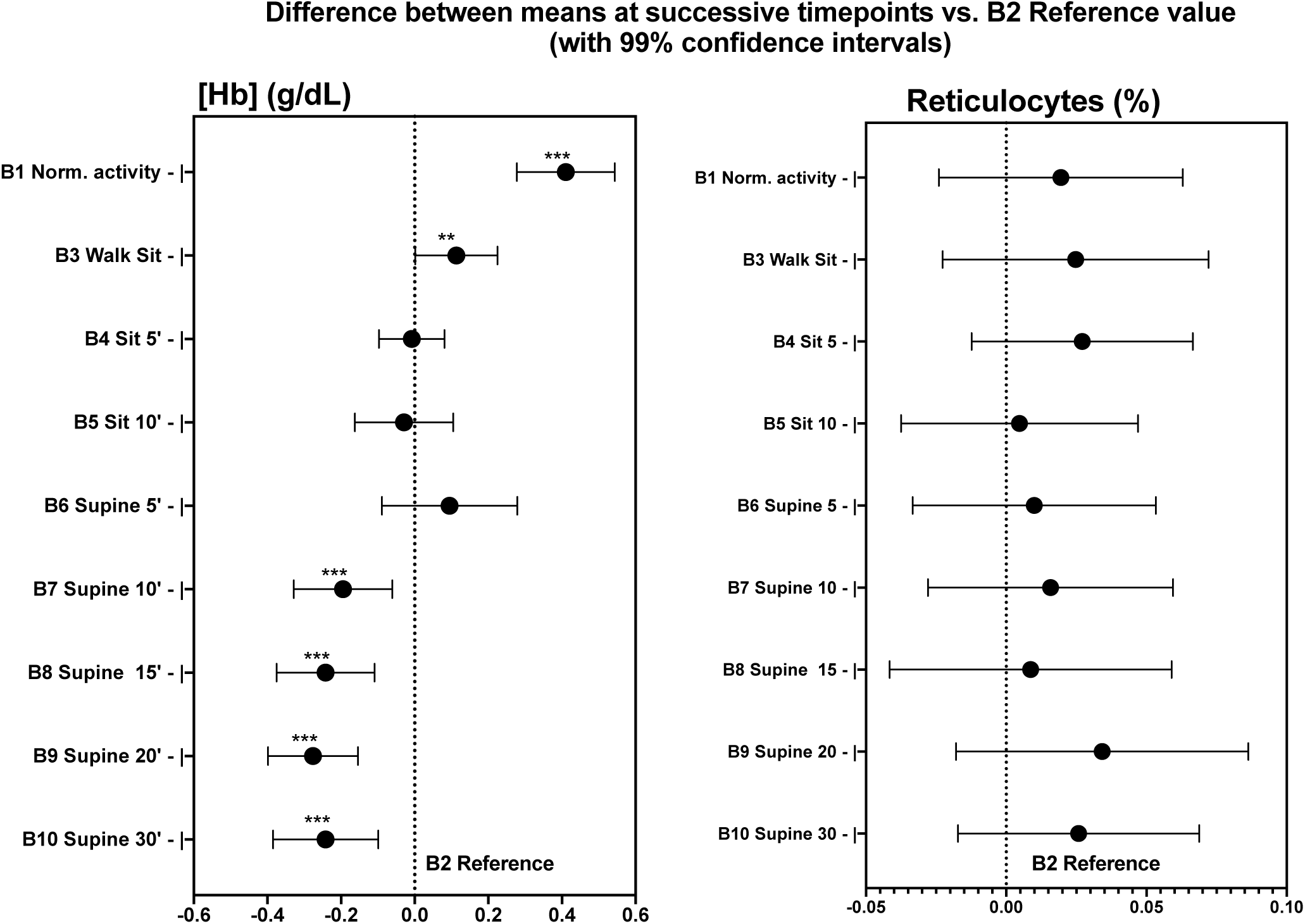
Difference between different timepoints and B2 reference values for hemoglobin concentration and reticulocytes percentage with 99% confidence intervals. *** *P*< 0.001, ** *P* <0.01 for difference with baseline (B2 reference)

Plasma volume changes compared to B2 ranged from – 5.3% to 5.1% % with a significant decrease of - 1.4% in average (*P*<0.05) at B3, and with a mean increase of 2.3%, 3.3%, 3.7%, 3.0% at B7, B8, B9 and B10, respectively (*P*< 0.001).

There was a Group effect on [Hb] (*P*<0.05) indicating that the cyclists had higher values when compared to the apnea divers and controls; and similarly, an interaction of Sex on [Hb] and Hct (*P*<0.001) indicating the lower values measured in female subjects. However, no interaction (Group x measure or Sex x measure) was identified for the posture related changes of [Hb], Hct or ΔPv over the 10 timepoints in our study (NS) (Table 2).

**Table 2.** Statistical results for the influence of the various fixed effects (measure, group, gender) and interactions on hemoglobin concentration ([Hb]), hematocrit (Hct) and calculated plasma volume changes (ΔPV).

| Effect | [Hb] |  | Hct |  | ΔPV |  |
| --- | --- | --- | --- | --- | --- | --- |
|  | F | P | F | P | F | P |
| Measure | 42.083 | <0.001 | 38.073 | <0.001 | 43.624 | <0.001 |
| Sex | 39.216 | <0.001 | 32.703 | <0.001 | 1.17 | 0.28 |
| Group | 3.508 | 0.042 | 2.296 | 0.117 | 1.57 | 0.22 |
| Measure * Group | 0.426 | 0.982 | 0.516 | 0.95 | 0.41 | 0.98 |
| Measure * Sex | 0.951 | 0.481 | 1.403 | 0.186 | 1.31 | 0.22 |

## Discussion

The major finding of the present study is that standing up and walking 50 m after being seated for 10 minutes increased [Hb] and Hct, while values were not different from baseline after 5 minutes in a seated position again. The present study also underlined that blood samples collected after 10-30 min in supine position resulted in lower [Hb] and Hct values compared to a sample taken after 10 min seated.

In an antidoping context, our results indicate that digressing from current guidelines imposing 10 min seated before sampling may result in altered results for variables measured as concentrations, due to plasma volume shifts. Practically, however, if an athlete has to change seats (e.g., walking a few meters), only 5 min may be required seated again before collecting a sample acceptable for the ABP. Our results are in line with the initial study by Ahlgrim *et al*. showing that posture significantly influences blood volume responses in its volumes distributions (Ahlgrim, Pottgiesser et al. 2010). It may be hypothesized that a rapid pooling of blood to the lower extremities is due to the law of gravity (Jacob, Raj et al. 2005). This phenomenon has been largely documented with an increase in vascular transmural hydrostatic force (Lundvall and Bjerkhoel 1994). Our results are also in accordance with an older study where a drop in blood volume up to 8% from the seated to the standing position, and an increase up to 12% in supine position were reported (Hinghofer-Szalkay and Greenleaf 1987). An increased capillary pressure in the feet (that can exceed 12kPa) causing an outward filtration was mentioned as a possible explanation. The smaller change observed at B3 in the present study could be explained by the very short standing (and walking) time of 50 s. Our result however indicate that values rapidly stabilize again (i.e. after 5 min) when returning to the seated position after the brief walk.

In comparison with a seated position, a supine position induces a decrease in vascular resistance and a shift of approximately 500-1000ml of blood, representing approximately 12% of blood volume (Robertson 2008). This important shift may explain the hemodilution observed in our results in the supine position. Moreover, our study results also highlight the time course of hemodilution from the standing to the supine position, in line with research realized on patients undergoing hemodialysis (Inagaki, Kuroda et al. 2001): A hemodilution was apparent after 10 minutes in supine position as highlighted mainly by the lowered [Hb] and an increase in plasma volume. In a medical context, a study showed that a 4% increase in Hct values from supine to standing position with a PV decrease between 6 and 25 % (Jacob, Raj et al. 2005). The magnitude of changes in PV observed in our study compared to the seated phase is in line with previous research with variations ranging from -5% to +5%, showing interindividual variability (Ahlgrim, Pottgiesser et al. 2010). In other research, taller subjects had a higher PV volume decrease when standing up that the authors explained by a greater orthostatic load (Lundvall and Bjerkhoel 1994). However, our results with the significant hemoconcentration observed at B3 in our (significantly taller) Cyc group at B3 did not confirm the latter. The similar variations in [Hb] and Hct indicate that the initial circulating volume and total haemoglobin content does not influence variations due to posture changes (i.e. no group x measure variation).

Our study included almost one third of female subjects and our results indicate that sex had an effect on [Hb] and Hct (p < 0.001) with lower values at baseline that are well documented (Murphy 2014). However, the effect of sex on the changes in blood variables are not significant indicating that our conclusions may apply to both male and female subjects or independently of the initial [Hb] or Hct. As pre-analytical conditions are crucial for the quality of blood analyses, beyond postural changes, other factors have to be considered to ensure the quality of the measurements. For instance, tourniquet time was less than 20 seconds on average and the collection of the blood sample was completed in 45 to 60 s. Prolonged tourniquet use (> 180 s) was thus avoided since it may lead to alterations in [Hb] and Hct values due to the increase in venous pressure causing a fluid shift into the extravascular space (Kuipers, Brouwer et al. 2005). The stable Ret% values obtained over the 10 successive measurements can be interpreted as a positive quality indicator for both sample collection and analytical measurements with the automated analyzer (Sysmex XN1000) (Lombardi, Lanteri et al. 2011). The Ret% values in our study are in line with references values in the general population varying between 0.5% and 2.5% (Banfi 2008). Ret% are not modified by plasma volume change as they are measured as an absolute value.

In conclusion, our study indicates that standing up shortly during an antidoping blood collection process (and walking up to 50 m to change seats for example) significantly alters the [Hb] and Hct values in athletes and healthy subjects. Values however stabilized after 5 min upon returning in a seated position. Blood sampled in a supine position may result in lower [Hb] and Hct values that can affect an ABP profile. Blood samples for anti-doping purposes in the context of the ABP should therefore not be collected in a supine position. If a subject has to stand up shortly after having waited for 10min (e.g., to change seat from a waiting room to the phlebotomy location), acceptable samples could be obtained after 5 min in a seated position. These findings can complement the current WADA guidelines for blood sampling in the context of the ABP.

## Acknowledgements

The authors wish to acknowledge WADA’s Science Department for the financial support of this study and all the participants for their participation.

## Author Contributions Statement

RF, YOS, MS and TA conceived the project. RF and MS obtained the project funding. TA contributed to the collection of data. TA, RF and FCvR statistically analyzed the data. TA, RF, YOS and MS drafted the final version of the manuscript. All authors contributed to revising the manuscript and expressed their approval of the final submitted version.

